# Auditory prediction errors as individual biomarkers of schizophrenia

**DOI:** 10.1101/104547

**Authors:** J.A. Taylor, N. Matthews, P.T. Michie, M.J. Rosa, M.I. Garrido

## Abstract

Schizophrenia is a complex psychiatric disorder, typically diagnosed through symptomatic evidence collected through patient interview. We aim to develop an objective biologically-based computational tool which aids diagnosis and relies on accessible imaging technologies such as electroencephalography (EEG). To achieve this, we used machine learning techniques and a combination of paradigms designed to elicit prediction errors or Mismatch Negativity (MMN) responses. MMN, an EEG component elicited by unpredictable changes in sequences of auditory stimuli, has previously been shown to be reduced in people with schizophrenia and this is arguably one of the most reproducible neurophysiological markers of schizophrenia.

EEG data were acquired from 21 patients with schizophrenia and 22 healthy controls whilst they listened to three auditory oddball paradigms comprising sequences of tones which deviated in 10% of trials from regularly occurring standard tones. Deviant tones shared the same properties as standard tones, except for one physical aspect: 1) duration-the deviant stimulus was twice the duration of the standard; 2) monaural gap-deviants had a silent interval omitted from the standard, or 3) inter-aural timing difference, which caused the deviant location to be perceived as 90° away from the standards.

We used multivariate pattern analysis, a machine learning technique implemented in the Pattern Recognition for Neuroimaging Toolbox (PRoNTo) to classify images generated through statistical parametric mapping (SPM) of spatiotemporal EEG data, i.e. event-related potentials measured on the two-dimensional surface of the scalp over time. Using support vector machine (SVM) and Gaussian processes classifiers (GPC), we were able classify individual patients and controls with balanced accuracies of up to 80.48% (*p*-values = 0.0326, FDR corrected) and an ROC analysis yielding an AUC of 0.87. Crucially, a GPC regression revealed that MMN predicted global assessment of functioning (GAF) scores (correlation = 0.73, R^2^ = 0.53, *p* = 0.0006)

## 1. Introduction

Schizophrenia is a chronic psychiatric disorder affecting approximately 1% of the population, expressed through cognitive dysfunction and psychotic symptoms such as hallucinations and delusions (Kahn et al., 2015). Schizophrenia is currently diagnosed through symptomatic evidence collected through patient interview. An investigation of current International Classification of Diseases diagnostic criteria (ICD-10, codes F20.0-F20.3 and F20.9) suggests the validity of schizophrenia diagnoses may be of about 89.7% (Uggerby et al., 2013). Whilst reasonably accurate, this method relies on self-report measures and ultimately on a subjective clinical decision. Hence, there is a pressing need to find biomarkers for schizophrenia that can objectively inform diagnosis and prognosis.

A number of potential candidates for schizophrenia biomarkers have been investigated, with the mismatch negativity (MMN), being one of them. The MMN is an event-related potential (ERP) elicited by an occasional unpredicted change (or deviant) in a sequence of predicted auditory events (standards). Indeed, the MMN is known to be robustly attenuated in patients with schizophrenia (Catts et al., 1995; Shelley et al., 1991; Todd et al., 2013) and is correlated with poor cognitive function (Light and Braff, 2005). MMN reduction is arguably one of the most reproducible neurophysiological markers of schizophrenia (Kaser et al., 2013; Shelley et al., 1991). Remarkably, this reduction is accentuated in people at risk who end up developing schizophrenia, compared to those who do not, even if there are no other behavioural differences at the baseline (Bodatsch et al., 2011; Perez et al., 2014).

While these are exciting findings, they rely on the comparison of group differences. In recent years, however, machine learning has been applied to neuroimaging data in order to provide predictive measures of diagnostic outcomes at the single individual level (Iwabuchi et al., 2013). For example, Gould et al. (2014) used abnormalities found in the neuroanatomical structure through MRI to classify schizophrenia patients and healthy controls with up to 72% accuracy.

Previous studies on EEG-based classification of schizophrenia via machine learning have primarily coupled auditory components of the ERP with visual attentional measures as discriminatory features. Neuhaus et al. (2014) measured the visual P300 response to unexpected sequences of letters, and the auditory P300 and MMN responses to a frequency oddball paradigm. In that study, they achieved an accuracy of 72.4% using the standard visual response at two electrode locations and nearest neighbour classification. Similarly, Laton et al. (2014) measured the visual P300 response to sequences of shapes, as well as the auditory P300 and MMN responses to a combined frequency and duration stimulus paradigm, achieving accuracies of up to 84.7% for a combination of all three paradigms, and up to 75% for MMN alone. However, it appears that these accuracy levels may potentially be inflated through the use of both model training and testing data for feature selection, rather than the training dataset alone. This and other studies, such as Neuhaus et al. (2011), are also limited to specific peak components (MMN and P300) as features, extracted through pre-processing in predefined time windows from discrete electrode locations.

The aim for this study was to develop a computational model which aids schizophrenia diagnosis, based on objective biological quantities, measured through widely accessible imaging technologies such as EEG, and a simple task suitable for patients. Instead of using predefined time windows and electrodes, we used a whole spatiotemporal approach by considering all the electrodes and the whole peristimulus window as potential features. We assessed the performance of multivariate pattern recognition in classification of schizophrenia patients and healthy controls, using three different auditory oddball paradigms. Moreover, we compared the performance of different classification algorithms (SVM vs. GPC), responses (standards, deviants, and MMN difference wave), feature selection (with and without an *a priori* defined temporal mask), and data normalisation operations.

## Methods

### Participants

Twenty-one individuals with schizophrenia (age 20-52 years, *M* = 39.7 years, *SD* = 9.0, 15 male) were recruited from outpatient sources, including a volunteer register managed by the Schizophrenia Research Institute and the Inner North Brisbane Mental Health Services of the Royal Brisbane Hospital. A healthy comparison group (*N* = 22) was recruited from students of the University of Newcastle and community volunteers. Control participants were similar to the schizophrenia patients in both age and sex (age 23-53 years, *M* = 39.1 years, *SD* = 9.4, 14 male). Controls recruited from the University of Newcastle received course credit for participation; all other participants were reimbursed for travel costs and expenses.

### Cognitive and clinical characterisation

Pre-morbid verbal IQ differed significantly between control (*M* = 117.5, *SD* = 6.8) and patient (*M* = 110.2, *SD* = 10.2) groups (*p* = 0.0078), based on the National Adult Reading Test (NART, Nelson, 1982). All participants were right handed, as assessed by the Edinburgh handedness inventory (Oldfield, 1971).

Participants were excluded if screening revealed a history of major head injury, epilepsy, hearing loss, or a recent history of substance abuse. Additionally, healthy controls were excluded if there was a personal history of mental illness, or a history of schizophrenia in first-degree relatives. Audiometric testing confirmed that detection thresholds were normal for all participants across frequencies of 500-2000Hz.

Diagnoses for individuals within the patient group were made using Diagnostic Interview for Psychosis (DIP, Castle et al., 2006). The same interview was administered to the healthy comparison group to exclude significant psychopathy. All patients included in this study received an ICD-10 diagnosis within the schizophrenia spectrum. Ratings of current symptomatology for patients were obtained on the Scale for Assessment of Positive Symptoms (SAPS, Andreasen, 1984) and the Scale for the Assessment of Negative Symptoms (SANS, Andreasen, 1982), summarised in Table 1. All patients were prescribed typical antipsychotic medication at the time of testing, except for one participant who was not receiving medication.

**Table 1.** Summary of schizophrenia patient symptom scores. Table shows group means and standard deviation for each measure of the SAPS and SANS (absent to severe, scale 0-5), and GAF (extremely high to severely impaired function, scale 1-100).

|  | Measure | Mean | SD |
| --- | --- | --- | --- |
| SAPS | Hallucinations | 2.20 | 1.74 |
|  | Delusions | 2.10 | 1.71 |
|  | Bizarre Behaviour | 1.05 | 1.19 |
|  | Positive Formal Thought Disorder | 0.75 | 1.16 |
|  | Summary SAPS Score | 6.10 | 3.92 |
| SANS | Affective Flattening | 2.00 | 1.17 |
|  | Alogia | 1.35 | 1.27 |
|  | Avolition | 2.55 | 1.00 |
|  | Anhedonia | 2.40 | 1.05 |
|  | Attention | 1.95 | 1.23 |
|  | Summary SANS Score | 10.25 | 4.67 |
| GAF |  | 55.43 | 14.93 |

The participants’ overall level of functioning, across psychological, social and occupational domains, was assessed using the Global Assessment of Functioning Scale (GAF, American Psychiatric Association, 2000), a numeric scale scored from 1-100 and divided into 10 associated levels of functioning and symptom severity. The GAF ratings ranged from 32-85 (*M* = 55.43, *SD* = 14.93) in patients and 73-90 (*M* = 83.8, *SD* = 5.3) in healthy controls.

### Experimental design

In an encephalographic auditory-oddball experiment, participants listened to sequences of short audio stimuli repeating at 500 ms intervals presented via headphones whilst watching a silent movie. Three different stimulus variations (Figure 1), each with specific tonal properties, were tested in separate blocks. For each paradigm, approximately a small percentage trials deviated from the standard stimulus in some physical aspect (8% for duration, 12% for left and right gap, and 10% for left and right inter-aural time difference deviants), occurring in a pseudo-random, non-consecutive order. These deviations were all expected to elicit the MMN signal.

**Figure 1.**
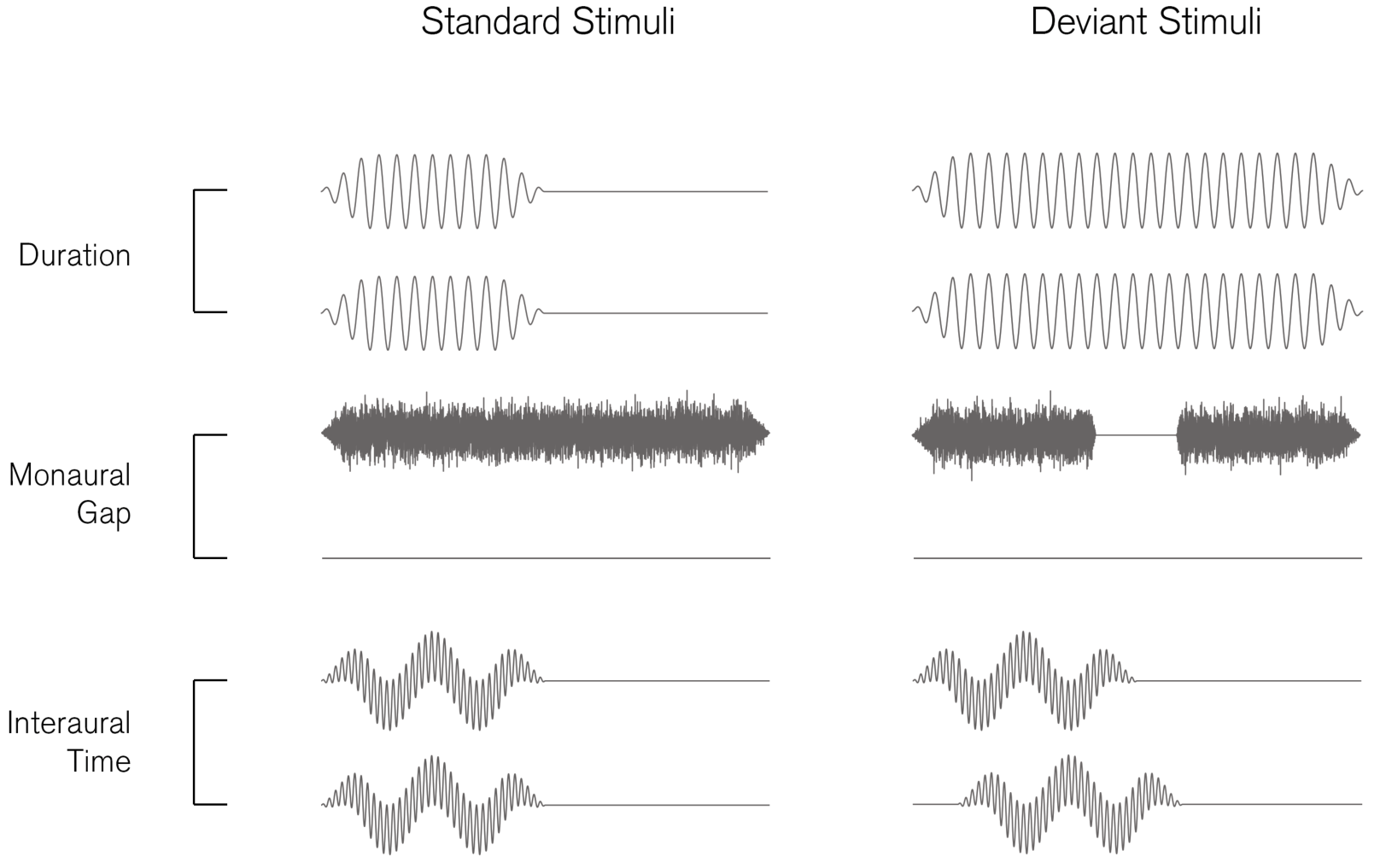
Binaural audio channel waveforms for standard and deviant stimuli used in each paradigm — duration (binaural 1 kHz sinusoidal tone with 50 ms standard duration, 100 ms deviant duration), monaural gap (white noise burst with 100 ms duration, deviant has 18 ms silent interval) and interaural time difference (6 kHz and 300 Hz harmonic tones with 50 ms duration, deviant has 0.7 ms delay between channels). *Note:* Waveforms are exaggerated for illustration and do not share common axes.

The first paradigm employed duration deviants (DUR, Figure 1a), where standard stimuli were binaural 1 kHz sinusoidal tones, 50 ms in duration, with deviant stimuli lasting 100 ms; i.e. twice the standard duration. The second was a gap paradigm (Figure 1b), for which standard stimuli were monaural bursts of white noise, 100 ms in duration, whereas the deviant stimuli contained an 18 ms silent interval centered within the noise. This paradigm was used for both left (GPL) and right ears (GPR) in separate blocks of the experiment. The third was an inter-aural time difference paradigm (ITD, Figure 1c), where standard stimuli comprised 6kHz and 300Hz harmonic tones, 50ms in duration. By playing identical tones simultaneously in both ears, the source location of standard stimuli was perceived to be central. Two oddball variations were used, such that the source of deviant stimuli was perceived to be located 90° either left or right of the midline. This effect was achieved through a 0.7 ms delay in one ear, creating a phase shift between the two ears. All stimuli were played at an 80 dB intensity level and were tapered using 10 ms rise/fall times for the duration and interaural time paradigms, 5 ms for rise/fall times for the gap paradigm and 1 ms transition in the corresponding gap deviant. The presentation of MMN paradigms was counterbalanced across participants. Given that the interaural time difference paradigm comprises a two binaural deviant stimuli and a common standard stimulus, these left and right deviant responses were averaged together, whereas the monaural gap paradigm comprises discrete standard and deviant stimuli pairs pertaining to the left or right channels in isolation. We therefore consider the interaural time to be a single paradigm, and the left and right gap to be two distinct variations of the same paradigm.

### Data collection and pre-processing

30 EEG sites (plus EOG) were collected using a 64-electrode head-cap at a sampling rate of 500Hz with referencing to the nose electrode. Offline signal processing was performed using the statistical parametric mapping toolbox (SPM12, Litvak et al., 2011). A band-pass Butterworth filter was applied with cutoff frequencies at 0.1 and 30 Hz. Experimental trials were then epoched with −100 to + 450 ms peri-stimulus intervals and baseline correction was applied using the -100 to 0 ms pre-stimulus period. Trials with signal amplitudes exceeding a ±100 µV threshold were excluded from the analysis. After artefact rejection, the mean number of stimuli analysed across all participants was 1222.2 standards and 153.5 deviants for the duration paradigm (13% of deviants), 712.7 standards and 112.8 deviants for the left gap (16% of deviants), 712.3 standards and 112.3 deviants for the right gap (16% of deviants), and 677.0 standards and 225.5 deviants for interaural time difference (33% deviants). Left gap paradigm data were excluded from one control participant due to an excessive number of trails containing these artefacts, whilst one patient was only subject to two of the three stimulus paradigms tested.

### Image conversion and feature definition

In this study, the averaged ERPs to standards, deviants, and MMN responses for each participant and stimulus paradigm were converted to a set of three-dimensional NIfTI images via SPM12. These images represent the two-dimensional surface of the scalp as a 32 **×** 32 matrix over 276 time points, with a sampling period of 2 ms. The resulting three-dimensional spatiotemporal volume forms the feature vector for each respective stimulus paradigm and response that is fed into the classifier. Additional masking was applied over either the 0-450 ms time interval, i.e. the full post-stimulus epoch period, or 50-250 ms, which was assumed to contain the auditory MMN component.

### Classification techniques

For classification of the control and patient groups, class labels were assigned to each subject. Using the Pattern Recognition for Neuroimaging Toolbox (PRoNTo), developed by Schrouff et al. (2013), we then applied two machine learning algorithms; Support Vector Machine and binary Gaussian Process Classifier. Support vector machines (SVM; Cortes and Vapnik, 1995) and Gaussian process classifiers (GPC; Rasmussen, 2006) are two of the most commonly used machine learning techniques in neuroimaging.

In the SVM training phase, weights are assigned to these features for maximal separation between the groups using a hyperplane, which serves as the decision boundary. Classification labels are determined by the sign of the total features weights multiplied by test sample. We use the default soft-margin parameter of C=1. GPC use probabilistic modelling to estimate the likelihood that a test sample belongs to a particular group. Given the covariance between samples in the training set and observed properties of the test sample, probability distributions of group membership are used to make predictions and assign a group label which best explains the data.

To assess the performance of models generated by these algorithms, we employed a *k*-fold cross-validation scheme, which divides the collective group of subjects into *k* subgroups or ‘folds’. These subgroups are iteratively assigned for training or testing the model until all subgroups have been used for testing. In this case, 10-folds (*k* = 10) cross-validation was used, where 90% of the data is used as the training set and the remaining 10% is used for testing the model. The number of subjects from both control and patient groups were balanced within each fold. This classification process also comprised additional data operations applied to both training and testing sets within the cross-validation loop. Mean-centering was applied to all models, in which the voxel-wise mean is subtracted from each data vector, and were computed both with and without normalisation, whereby data vectors are divided by their Euclidian norm. In a neuroimaging context with small sample sizes such as these, leave-one-out cross-validation is commonly applied (Veronese et al., 2013). However, by taking a more conservative approach using 10-folds in this manner we hope to provide a more robust measure of performance. Finally, to determine the statistical significance of the results, permutation tests were calculated for each model and cross-validation scheme, retraining the model with randomised labels and 1000 repetitions.

When comparing models generated with all possible combinations of the four stimulus paradigm variations, three responses, two time intervals, two data operations and two algorithms presenting in this study, we arrive at a total 96 hypotheses. To correct for multiple comparisons, we applied false discovery rate controlling procedures (Benjamini and Hochberg, 1995; Benjamini and Yekutieli, 2001) with a desired false discovery rate of *q* = 0.05. Assuming patient diagnoses in the dataset were accurate, we therefore do not consider above chance accuracies to be positive results unless a critical uncorrected significance level of *p* ≤ 0.005 is achieved via permutation testing. These permutation tests were repeated using 10,000 repetitions for models which met this condition.

### Regression modelling

To investigate quantifiable relationships between neuronal responses and symptoms, the control and schizophrenia patient groups were combined into a single group and spatiotemporal images were mapped to their available SANS, SAPS and GAF diagnostic scores. Within the PRoNTo toolbox, three standard multivariate regression algorithms were applied; Kernel Ridge Regression (KRR; Shawe-Taylor and Cristianini, 2004), Relevance Vector Regression (RVR; Tipping, 2001) and Gaussian Process Regression (GPR; Rasmussen and Williams, 2006).

KRR derives a relationship between samples by minimising an error function comprising the sum of the squared differences between model predictions and regression targets and a model regularization term, which in turn reduces the size of weights and possible overfitting. RVR is similar to SVM, however uses conditional probability to make estimations rather than absolute predictions. The model weighting ‘vectors’ are initially assigned a Gaussian prior with mean zero, which are then iteratively optimised via the training process, after which those with non-zero means are considered the most ‘relevant’ in making predictions. GPR use the same methodology as GPC.

Again, the performance of these models is assessed using 10-fold cross-validation and the statistical significance of results is assessed via permutation testing with Sidák correction for multiple comparisons.

## Results

### Group responses

The standard and deviant responses collected from the CZ channel, coupled with the MMN difference waves are plotted for all paradigms in Figure 2a, with individuals shown as dotted lines alongside group averages (schizophrenia in colour and control in greyscale). Patients with schizophrenia displayed higher variability in the responses to both standard and deviant conditions.

**Figure 2.**
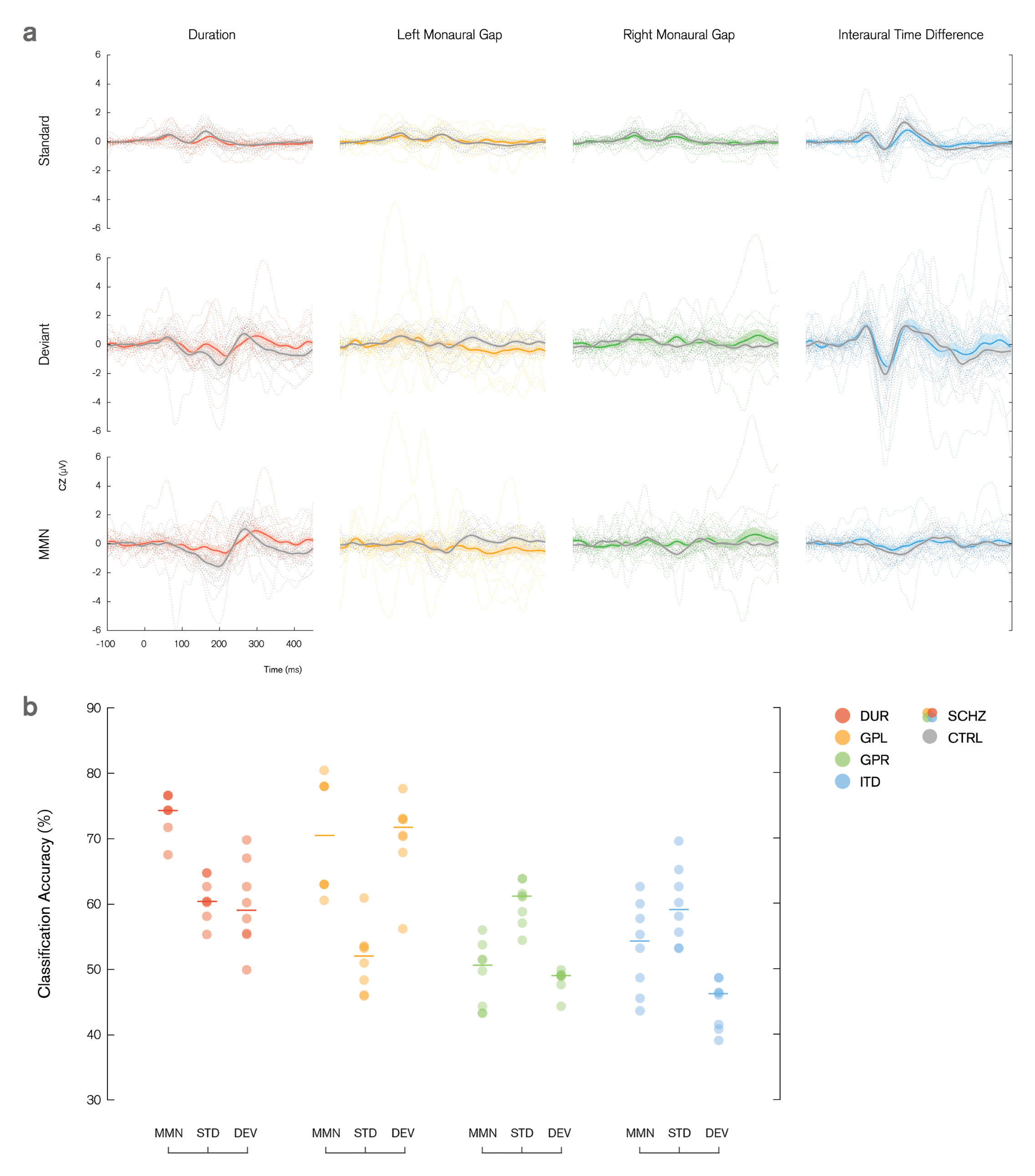
***pictured over* (a)** Averaged standard, deviant and MMN responses to duration (red), left gap (yellow), right gap (green) and interaural time difference (blue) stimulus paradigms, recorded from the CZ channel for control group (grey) and schizophrenia patients (colour). Individual subjects are denoted using dotted lines. Shading illustrates the standard error of the mean. **(b)** Balanced classification accuracy distributions for each stimulus paradigm with respect to measured response (MMN, standard and deviant). The number of samples with a particular accuracy level are represented with increased colour intensity. Medians of each set are indicated using horizontal lines.

The MMN difference waves from the left gap paradigm performed best in our classification analysis, and can be examined in greater detail in Figure 3a (schizophrenia in yellow and control in grey). The magnitude of the difference wave averaged across the control group increases later in post-stimulus, peaking at approximately 250 ms, and remaining amplified thereafter. The average difference wave in the schizophrenia group response remains fairly flat for the full epoch length. This effect is further emphasised through normalisation of the ERPs, as illustrated in Figure 3b.

**Figure 3.**
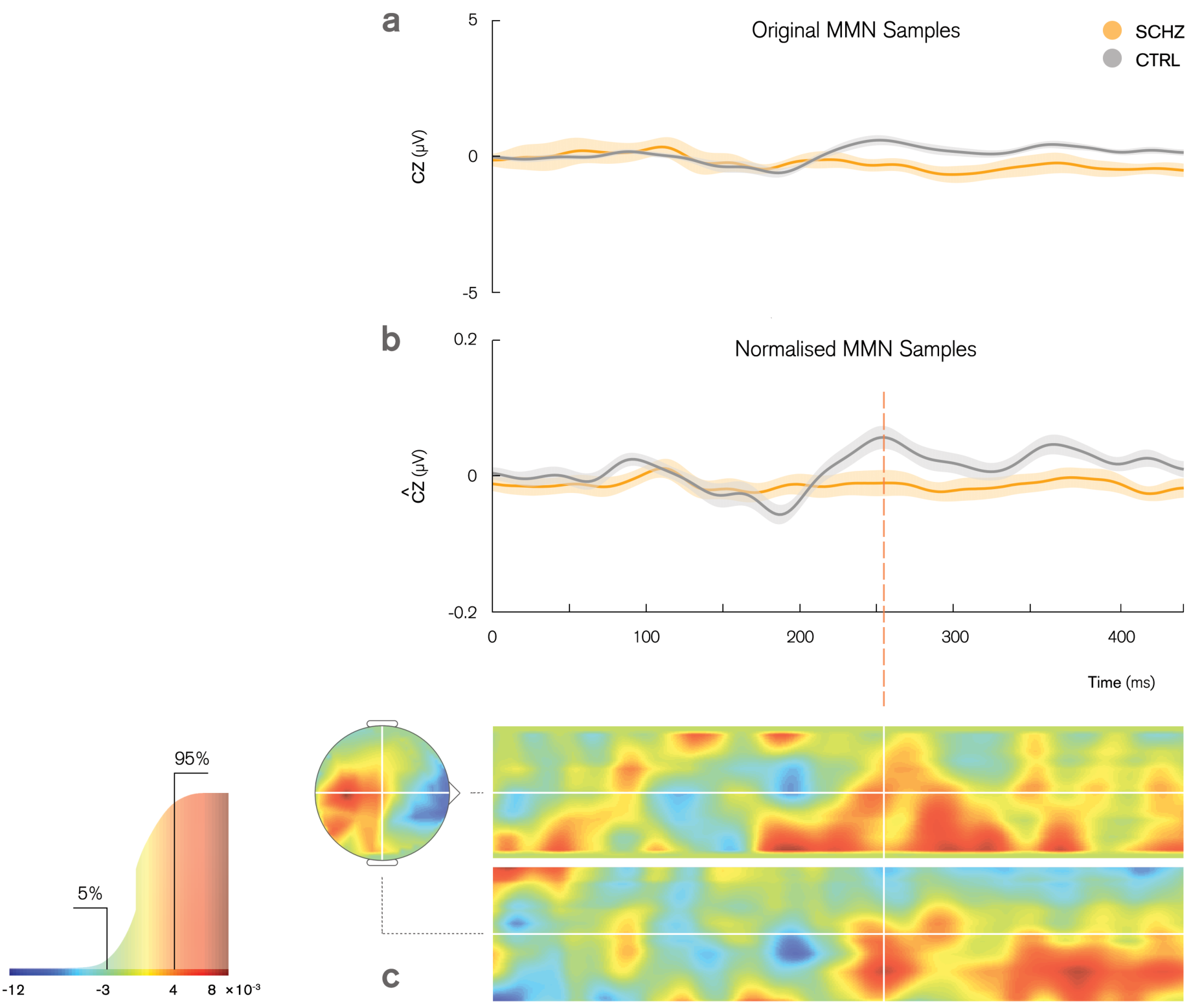
**(a)** Left gap MMN recorded from CZ channel for control (grey) and schizophreniagroups (yellow) with grand means and S. E. M. across groups. **(b)** Effect of normalisation on the MMN signal. **(c)** Sample weights map showing distribution of weights assigned to spatiotemporal features. Associated colour mapping is shown on the cumulative distribution function graph (left), indicating the 5% most positive (red) and 5% most negative (blue) weights.

The points in space and time which contributed most to model decision making can be identified through the weights map which achieved highest classification accuracy. Weights are parameters obtained through the initial training phase, representing the relative contribution of each voxel to the classification decision. In the testing phase, the classifier will compute the product of a given spatiotemporal sample and this weight mapping to obtain a function value, which is then used to make the prediction of group membership.

The weight map is shown in Figure 3c, with spatial guides projected through the location of the Cz channel on the scalp, as indicated in white. The top and bottom 5% of weights displayed in this figure (values less than −3 × 10^-3^ and greater than 4 × 10^-3^) are highlighted for visualisation purposes in the cumulative distribution of all signed weights. Note however, that all voxels, including those with low assigned weights, contribute to the prediction. As labeled on the colour scale, positive and negative weights are shown in red and blue respectively. A positive weight applied to a higher ERP amplitude at a particular point in space and time produces a larger function value, contributing to a healthy control classification. In this instance, large clusters of positive weights have been assigned to the mismatch response amplitude from 250ms onward, indicating that the algorithm has detected these time points as important in discriminating between the two groups, although individual weights should only be interpreted within the context of the full weight map.

### Analysis of model performance

The performance of prospective models is measured through total and balanced classification accuracies, class accuracy and class predictive value, calculated through the cross-validation testing phase. To assess the relative impacts of each parameter on model performance, the balanced accuracies from all models were partitioned into subsets according to the algorithm, paradigm, response, time-interval mask and normalisation operations applied. We first used the Shapiro-Wilk test to test for normality in each subset, using a significance level of **α** = 0.05. This hypothesis was rejected across all parameters. We were unable to normalise the data through any standard transformation due to the varied distributions across factors. Consequently, five non-parametric Kruskall-Wallis tests, were used to determine the main effects of algorithm (SVM, GPC), paradigm (DUR, GPL, GPR, ITD), response (standard, deviant, MMN), mask (0-450 ms, 50-250 ms), and normalisation (with, without) on classification accuracy. The results from these tests are shown below in Table 2, indicating a significant effect of paradigm. Contrary to our initial hypothesis, there was no marked improvement in classification accuracy when applying the 50-250 ms time interval mask as opposed to the full 0-450 ms epoch. No significant differences in overall performance were found between algorithms, responses or normalisation operations.

**Table 2.** Kruskall-Wallis tests comparing main effects of algorithms, paradigm, response, temporal masks and normalisation on balanced classification accuracy, demonstrating that paradigm manipulations have a significant effect. *p*-values are Sidák corrected for multiple comparisons.

| Sources | Sum of Squares | Degrees of Freedom | Mean Square | Chi-sq. | p > Chi-sq. Uncorrected | p > Chi-sq. Sidák Corr. |
| --- | --- | --- | --- | --- | --- | --- |
| Algorithm | 404.3 | 1 | 404.26 | 0.52 | 0.4704 | 0.9583 |
| Paradigm | 23522.4 | 3 | 7840.80 | 30.32 | **1.1813E-06** | **5.9065E-06** |
| Response | 3742.5 | 2 | 1871.26 | 4.82 | 0.0896 | 0.3746 |
| Mask | 2016.7 | 1 | 2016.67 | 2.60 | 0.1069 | 0.4318 |
| Normalisation | 437.8 | 1 | 437.76 | 0.56 | 0.4525 | 0.9508 |

The relative performance of different paradigms was then further examined through non-parametric right-tailed Wilcoxon rank-sum tests for pairwise comparisons (Table 3). The independent pairs of parameters compared in a rank-sum test are expressed in competition. The test itself calculates the probability that there is a difference in medians between these two data vectors, in this case whether the former is greater than the latter (e.g. standard *vs.* deviant, ‘is the median accuracy from the standard response greater than that from the deviant’?).

**Table 3.** Follow-up comparison of stimulus paradigms, and the effects of algorithms, measured responses, time interval masks and normalisation operations within the two best paradigms (duration and left gap) using Wilcoxon rank-sum test. *p*-values are Sidák corrected for multiple comparisons.

| Sources |  | Parameter Comparison | p-value | Rank |
| --- | --- | --- | --- | --- |
| Paradigm |  | DUR vs. ITD | **4.1775E-06** | 792.0 |
|  |  | DUR vs. GPL | 0.5020 | 588.0 |
|  |  | DUR vs. GPR | **9.9410E-07** | 803.5 |
|  |  | GPL vs. ITD | **2.2837E-04** | 753.0 |
|  |  | GPL vs. GPR | **2.4913E-04** | 752.0 |
|  |  | GPR vs. ITD | 0.3815 | 603.0 |
| DUR | Algorithm | SVM vs. GPC | 0.9839 | 147.5 |
|  | Response | MMN vs. Standard | **0.0004** | 100.0 |
|  |  | MMN vs. Deviant | **0.0008** | 99.0 |
|  |  | Deviant vs. Standard | 0.9988 | 62.0 |
|  | Mask | 0-450ms vs. 50-250ms | 0.3635 | 174.0 |
|  | Normalisation | None vs. Normalised | 0.5877 | 167.5 |
| GPL | Algorithm | SVM vs. GPC | 0.8650 | 158.0 |
|  | Response | MMN vs. Standard | **0.0008** | 99.0 |
|  |  | MMN vs. Deviant | 0.8843 | 72.0 |
|  |  | Deviant vs. Standard | **0.0008** | 99.0 |
|  | Mask | 0-450ms vs. 50-250ms | 0.3639 | 174.0 |
|  | Normalisation | None vs. Normalised | 0.9905 | 145.5 |

The *p*-values presented in Table 3 suggest the duration and left gap paradigms were comparable in performance, however both had significantly higher classification accuracy in contrast with the interaural time difference and right gap paradigms.

To follow this up, we then computed an equivalent interaction between paradigm and other factors within the best paradigms, as shown in Table 3. For the duration paradigm, MMN significantly outperformed individual standard and deviant responses, whereas for the left gap paradigm, MMN and deviant performed comparably with higher accuracy than the standard response. These findings are also illustrated in Figure 2b, which displays the accuracy of all classifiers with respect to the most significant factor of stimulus paradigm, split into the different types of responses. The number of models with a particular accuracy level are shown with increasing colour intensity, and median accuracy indicated by horizontal lines.

The models which survived multiple comparisons correction at the corrected alpha level of 0.05 are displayed in Table 4. The remaining models which did not survive correction for multiple comparisons are available in the appendix. The model with the best accuracy was obtained using the GPC algorithm with the MMN response to the left gap paradigm as features, further processed with the 0-450 ms temporal mask and normalisation operations.

**Table 4.** Summary of best performing classification models which survived multiple comparisons correction at the corrected alpha level of 0.05. Duration and left gap paradigms yielded the top balanced accuracies, ranging from 74.34% to 80.48% with FDR-corrected *p*-values above chance.

| Paradigm | Mask | Response | Algorithm | Normalisation | Total accuracy | Balanced accuracy | BA p-value uncorrected | BA p-value FDR corr. |
| --- | --- | --- | --- | --- | --- | --- | --- | --- |
| Duration | 0-450ms | MMN | GPC | None | 74.42% | **74.35%** | 0.0041 | **0.0437** |
|  | 50-250ms | MMN | SVM | None | 76.74% | **76.73%** | 0.0031 | **0.0372** |
|  |  |  |  | Normalised | 76.74% | **76.62%** | 0.0029 | **0.0372** |
|  |  |  | GPC | None | 76.74% | **76.62%** | 0.0025 | **0.0372** |
| Left Gap | 0-450ms | MMN | SVM | None | 78.05% | **77.98%** | 0.0010 | **0.0326** |
|  |  |  |  | Normalised | 78.05% | **77.98%** | 0.0017 | **0.0326** |
|  |  |  | GPC | None | 78.05% | **77.98%** | 0.0016 | **0.0326** |
|  |  |  |  | Normalised | 80.49% | **80.48%** | 0.0009 | **0.0326** |
|  | 50-250ms | Deviant | SVM | None | 78.05% | **77.74%** | 0.0017 | **0.0326** |

The histogram of returned function values from this model are shown in Figure 4a. These values are the probability of the subject belonging to the control group and are used to determine which labels are assigned to each subject, relative to the decision boundary at 0.5. In an ideal classification model, these two distributions would not overlap and while there is some overlap here, the model does a very decent job of separating the two groups.

**Figure 4.**
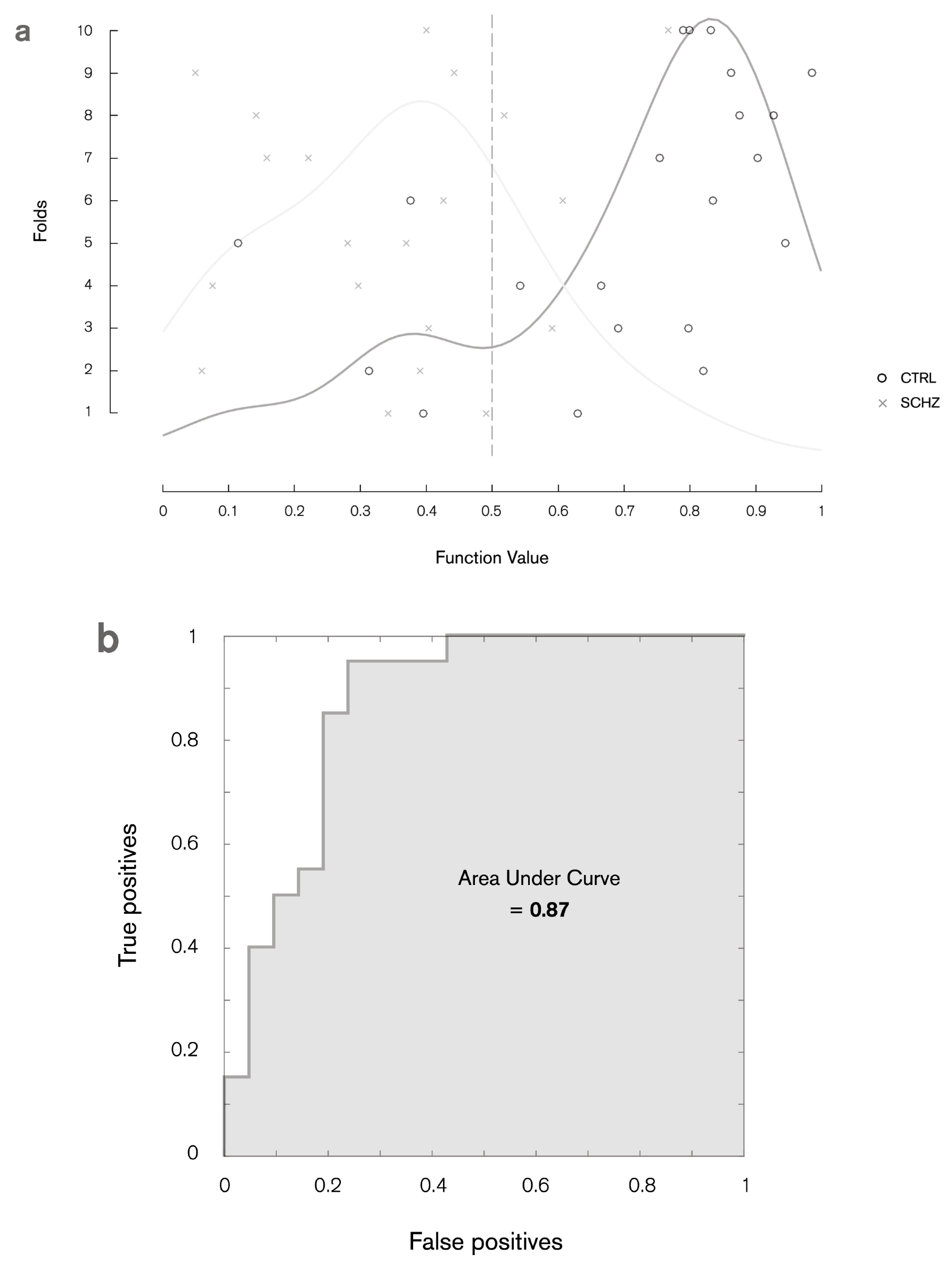
**(a)** Histogram of model function values and binary decision plot illustrating classification accuracy of individual subjects from controls (black circles) and schizophrenia patient groups (grey crosses). **(b)** ROC curve analysis with an AUC of 0.87. Values are displayed for the best classifier (80.49% accuracy, Gaussian Process Classifier using MMN features over 0-450 ms time window with normalisation operations).

Model sensitivity and specificity is illustrated through the receiver operating characteristic (ROC) curve shown in Figure 4b, plotting the true positive rate (sensitivity) as a function of false positive rate (1 - model specificity). The area under the curve (AUC) measures how well the model distinguishes between subjects with and without schizophrenia.

These results informed exploratory regression modelling, in which the spatiotemporal images from each participant were mapped to their available diagnostic scores. The performance of these models is assessed by comparing the predicted and target values using Pearson’s correlation coefficient (*R*) and the coefficient of determination (*R*^2^, the ratio between the explained and total variance). Preliminary analyses of the SANS and SAPS from the schizophrenia patient group (*N* = 20) did not indicate any significant relationship, and of the four stimulus paradigm variations, only the left gap paradigm produced significant correlation with the GAF using the combined control and patient groups (*N* = 39). As shown in Table 5, these models performed consistently across KRR, RVR and GPR algorithms, and those which include the normalisation operation produced higher correlation (73-74%, *p* ≤ 0.0006, Sidák corrected) and explained variance (53-55%, *p* ≤ 0.0006) than those which did not (*R* = 0.59, *R*^2^ = 0.35, *p* ≤ 0.003). The GAF scores for two controls were not recorded at the time of experiment, and hence excluded from this analysis. The GAF scores predicted by the model which produced lowest mean square error (MSE = 156.08, *p* ≤ 0.0006) are plotted in Figure 5.

**Table 5.** Regression modelling performance measures using individual MMN responses to the left gap stimulus paradigm with 0-450 ms temporal mask, mapped to participants’ GAF scores. *p*-values are Sidák corrected for multiple comparisons.

| Algorithm | Normalisation | Correlation | R p-value<br>uncorrected | R p-value<br>Sidák corr. | R<sup>2</sup> | R<sup>2</sup> p-value<br>uncorrected | R<sup>2</sup> p-value<br>Sidák corr. | MSE | Norm.<br>MSE | MSE p-value<br>uncorrected | MSE p-value<br>Sidák corr. |
| --- | --- | --- | --- | --- | --- | --- | --- | --- | --- | --- | --- |
| Kernel Ridge<br>Regression | None | 0.59 | 0.0005 | 0.0030 | 0.35 | 0.0010 | 0.0060 | 342.42 | 5.90 | 0.0107 | 0.0625 |
|  | Normalised | 0.74 | 0.0001 | 0.0006 | 0.55 | 0.0001 | 0.0006 | 195.20 | 3.37 | 0.0001 | 0.0006 |
| Relevance Vector<br>Regression | None | 0.59 | 0.0002 | 0.0012 | 0.35 | 0.0006 | 0.0036 | 342.48 | 5.90 | 0.0371 | 0.2029 |
|  | Normalised | 0.73 | 0.0001 | 0.0006 | 0.53 | 0.0001 | 0.0006 | 156.08 | 2.69 | 0.0001 | 0.0006 |
| Gaussian Process<br>Regression | None | 0.59 | 0.0002 | 0.0012 | 0.35 | 0.0005 | 0.0030 | 342.43 | 5.90 | 0.0116 | 0.0676 |
|  | Normalised | 0.73 | 0.0001 | 0.0006 | 0.53 | 0.0001 | 0.0006 | 156.33 | 2.70 | 0.0001 | 0.0006 |

**Figure 5.**
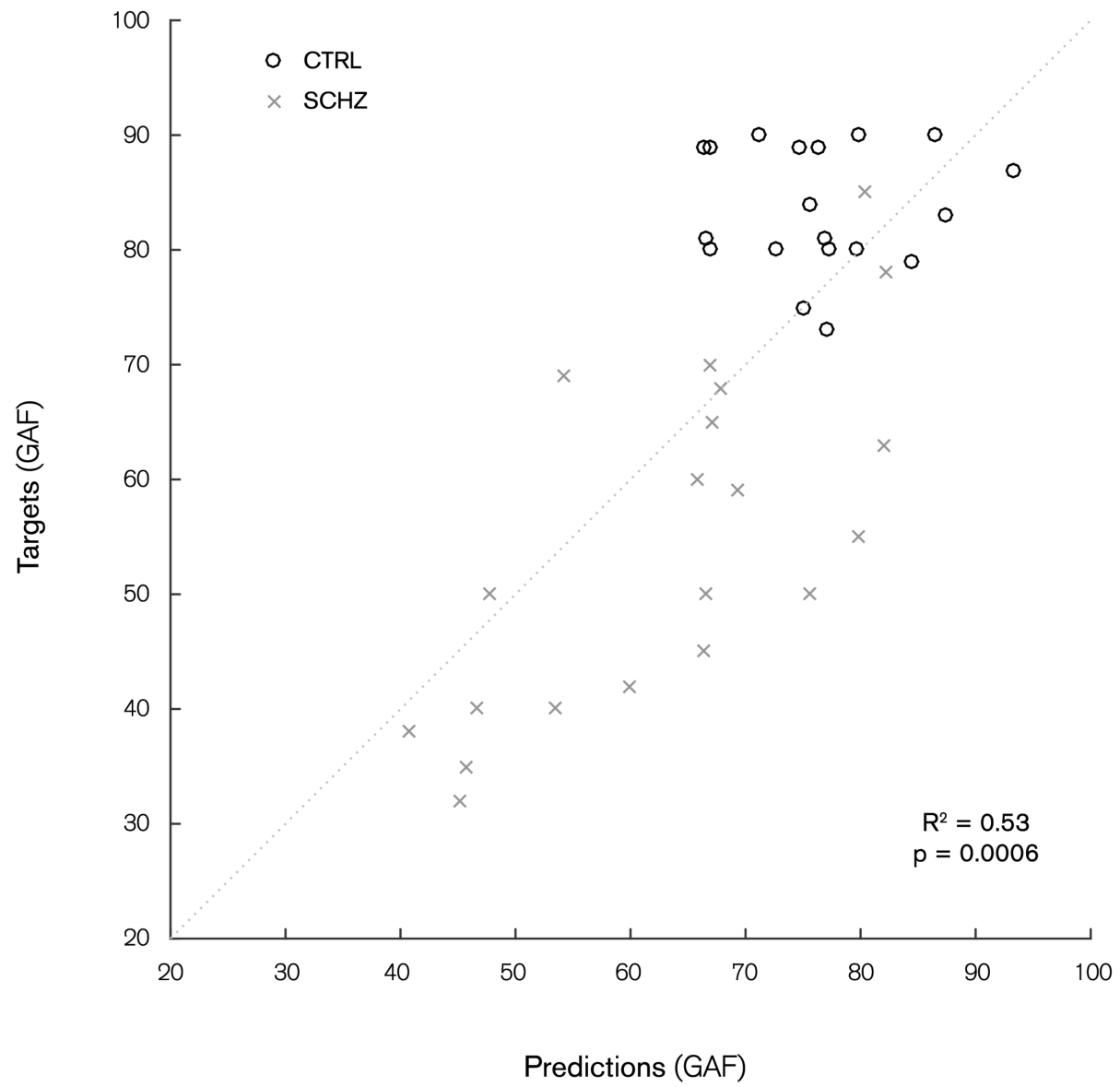
Plot of predicted and target GAF scores (scale 1-100). Individual subjects (black circles) and schizophrenia patients (grey crosses) are modelled as a single group. Values are assigned by Gaussian Process Regression using MMN features from the left gap paradigm over 0-450 ms time window with normalisation operations (73% correlation, p=0.0006 Sidak corrected, 53% explained variance, 156.08 mean square error).

## Discussion

This study investigated the application of multivariate pattern analysis (MVPA) techniques in the classification of people with schizophrenia and a matched healthy cohort. Participants listened to sequences of tones that changed occasionally in three different physical properties: sound duration, gap (silent interval within the tone), or perceived source location (through changes in interaural time). All of these oddball paradigms have been previously shown to elicit the MMN signal (Näätänen et al., 2004). Using spatiotemporal images as feature sets, Support Vector Machines and Gaussian Process Classifiers enabled us to classify individuals into their corresponding groups with up to 80.48% balanced accuracy (*p* ≤ 0.0326, FDR corrected). The most robust models presented here were generated through Gaussian process classification of MMN features, as measured over a 0-450 ms time window in response to the left gap stimulus paradigm. The regression model which had the smallest mean square error (relevance vector regression, normalised data) revealed 73% correlation (*p* ≤ 0.0006, Šidák corrected) between the left gap MMN response and available GAF scores, with explained variance of 53% (*p* ≤ 0.0006, Sidák corrected).

Although a number of similar schizophrenia classification studies have used machine learning algorithms on auditory oddball responses, this is the first to comparing several types of oddball paradigms. Moreover, it departs methodologically from different studies in that it takes the whole spatiotemporal information in the EEG data as potential features for the classifiers. Neuhaus et al. (2014) used a combined click-conditioning and auditory oddball paradigm with square wave standards and sinusoidal deviants, explicitly targeting the P50, N100 and P300 components. Our approach yielded a greater balanced classification accuracy (up to 80.48% in our study compared to 77.7%), and our ROC analysis indicated that the left gap paradigm provides greater discrimination between groups with an AUC of 0.87 outperforming 0.737 reported in (Neuhaus et al., 2014).

More direct comparisons can be made with a study by Laton et al. (2014), who targeted the MMN response directly with a combined duration and frequency stimulus paradigm, achieving an accuracy of up to 75.0%. Whilst other attention-related paradigms were employed to elicit auditory and visual P300 responses, we suggest that a simpler experimental setup and measures such as MMN difference wave, which are known to be more reproducible, are preferable in a diagnostic setting. We also note that the individual components which define this feature set were extracted using the grand averages of the dataset, which may also introduce some bias to the learning phase, and hence inflate accuracy. We are also more conservative in our reporting with significance calculated via permutation testing and correction for multiple comparisons.

Du et al. (2012) achieved accuracies of over 90% using functional magnetic resonance imaging (fMRI) to measure both resting state data and responses to a task-associated auditory oddball paradigm comprising frequency and randomised digital deviants. These results are likely to be inflated due to the use of leave-one-out cross-validation on the sample of 28 schizophrenia patients and 28 healthy controls. In comparison with fMRI, EEG is also able to achieve higher temporal resolution with simpler technology which is much more accessible to a wider population.

We used a 50-250 ms time interval mask, which was assumed *a priori* to contain the auditory N100 and MMN component, features which are considered to be the most reproducible neurophysiological markers of schizophrenia (Kaser et al., 2013; Shelley et al., 1991). We hypothesised that isolation of these features would maximise model performance, however in contrast, our best performing model resulted from the full 0-450 ms post-stimulus epoch feature set. Further region-of-interest weight map analysis indicated that the time points selected by the algorithm as most relevant in the classification were actually outside of the 50-250 ms interval, although all spatiotemporal voxels contribute globally to the prediction. The N100 and MMN are the most widely studied neurophysiological markers in schizophrenia, but abnormalities have also been reported in other event-related potential components, namely decreased attenuation of the P50 auditory evoked potential, which is considered a measure of neuronal inhibition (Källstrand et al., 2012). Reduced amplitude and increased latency of the P300 have also been consistently reported in schizophrenia patients and correlates with the degree of MMN reduction (Javitt et al. 1995; Ford et al., 2010; Brown et al., 2013).

A key limitation when performing binary classification of two groups is that, in general, schizophrenia cannot be described as a binary condition, comprising many levels of severity, differing symptoms and categorical subtypes. Similarly, healthy controls may experience mild schizotypal symptoms which do not warrant clinical diagnosis, but suggest a ‘continuum of psychosis’ (Oestreich et al., 2016). In keeping with this idea, in a study by Light and Braff (2005), links have previously been made between MMN and GAF scores in schizophrenia patients using individual EEG sites and a duration stimulus paradigm, with a correlation of 65% that accounted for 42% of the total variance. In our regression model testing with the left gap paradigm, we were able to improve on both of these performance measures. Note, differences in the GAF are not specific to schizophrenic attributes, but to the general global functioning which can be impaired in other psychiatric illnesses. Preliminary testing using SANS and SAPS diagnostic scores have not proven significant for this sample (data not shown).

It is worth mentioning that the schizophrenia patients were medicated over the course of this experiment, whereas the control group was not. These dosages are unknown, unfortunately, and thus we are unable to use them as a covariate in model training, or to verify whether it is uncorrelated with our classifier outputs. However, it is unlikely that the effects observed in this study are driven by a confound of drug, as MMN reduction is observed in both medicated and un-medicated patients (Catts et al., 2005), and such reduction is also present in unmedicated prodromals (Bodatsch et al., 2011; Sheikh et al., 2012). Another important difference between our two groups is the IQ score, which was significantly higher for the healthy controls compared to the patients with schizophrenia. This is a difficult problem to circumvent given the known strong correlation between low IQ and schizophrenia (Kendler et al., 2015).

In both classification and regression modelling, performance was shown to be highly dependent on the stimulus paradigm and measured response used as the feature set. However, when considering the varied levels of performance, perhaps the starkest result was the difference between the left and right gap paradigms, an observation which is consistent with previous findings showing that: 1) temporal processing is particularly impaired in schizophrenia (Michie, 2001) and 2) spatial asymmetries in the perception of sounds present in neurotypical individuals is lacking in patients with schizophrenia (Matthews et al., 2007). For example, in healthy participants, ERPs elicited in an auditory oddball task with frequency deviation are shown to be sensitive to which ear receives stimulation (left monaural, right monaural or binaural), with right lateralisation of scalp topography in response to left ear stimulus and bilateral distribution for right ear stimulus (Gilmore et al., 2009). This asymmetry is thought to be caused by an engagement of parieto-fronto-temporal pathways on the right hemisphere when the deviant stimulus occurs on the left side of space, and bilateral engagement of this pathway when the stimulus occurs on the right side (Dietz et al., 2014). In addition, even earlier on in the auditory pathway, this hemispheric asymmetry in healthy participants and the lack thereof in patients with schizophrenia can be caused by a number of reasons, including information corruption between hemispheres. Indeed, there are known differences in the inter-hemispheric transfer time when patients with schizophrenia and healthy controls are presented with monaural word stimuli (Henshall et al., 2012). In the left gap paradigm, stimuli presented to the left ear are preferentially processed in the right hemisphere, and would necessitate an interhemispheric transfer back to left hemisphere (which is most specialized for temporal processing). An alternative possibility for such asymmetry lies on the degree of grey-matter loss in the left temporal lobe, which correlates with MMN attenuation (Näätänen et al., 2004). Collectively, these results suggest that differences in brain connectivity and signal transfer between hemispheres may be emphasised when the participant listens to lateralised stimuli. The responses to such stimuli could form valuable discriminatory features in machine learning for schizophrenia diagnosis.

In conclusion, we used spatiotemporal images of event-related potentials evoked in response to varied auditory oddball stimuli as features, and were able to discriminate schizophrenia patients from healthy controls through machine learning algorithms. The MMN difference wave signal was found to produce the highest classification accuracy in comparison to individual standard and deviant responses. Evoked responses outside the auditory P100 and MMN were also found to be relevant discriminatory features. The greatest classification accuracy was achieved using a monaural gap stimulus paradigm in the left ear, with balanced classification accuracies up to 80.48% (*p*-values ≤ 0.0326, FDR corrected). The relationship between MMN responses to the left gap paradigm and the GAF score was also shown to have 73% correlation and explained 53% of the total variance (*p*-values ≤ 0.0006, Sidák corrected).

## Acknowledgements

This work was funded by a University of Queensland Early Career Researcher Grant (2013002373) and Fellowship (2016000071) to MIG. Data collection was funded by the National Health and Medical Research Council NHMRC: (Project Grant ID 209828) and was supported by the Schizophrenia Research Institute (SRI) and Hunter Medical Research Institute (HMRI). We thank Stanley Catts for helping with recruitment, Anderson Winkler and Anton Lord for discussions on statistical analysis, and Janaina Mourao-Miranda for discussions on machine learning methods.

